# Host “cleansing zone” at secondary contact: a new pattern in host-parasite population genetics

**DOI:** 10.1101/2020.02.28.969279

**Authors:** Jana Martinů, Jan Štefka, Anbu Poosakkannu, Václav Hypša

**Author notes:** Author for correspondence: Václav Hypša, Department of Parasitology, University of South Bohemia, České Budějovice, Czech Republic.

## Abstract

We introduce a new pattern of population genetic structure in a host-parasite system that can arise after secondary contact (SC) of previously isolated populations. Due to different generation time and therefore different tempo of molecular evolution the host and parasite populations reach different degrees of genetic differentiation during their separation (e.g. in refugia). Consequently, during the SC the host populations are able to re-establish a single panmictic population across the whole recolonized area, while the parasite populations stop their dispersal at the SC zone and create a narrow hybrid zone (HZ). From the host’s perspective, the parasite’s HZ functions on a microevolutionary scale as a “host-cleansing filter”: while passing from area A to area B, the hosts are rid of the area A parasites and acquire the area B parasites. We demonstrate this novel pattern on a model composed of *Apodemus* mice and *Polyplax* lice by comparing maternally inherited markers (complete mitochondrial genomes, and complete genomes of vertically transmitted symbiont *Legionella polyplacis*) with SNPs derived from the louse genomic data. We discuss circumstances which may lead to this pattern and possible reasons why it has been overlooked in the studies on host-parasite population genetics.

## Introduction

Population genetics within a host-parasite association are complex systems dependent on many ecological features of both counterparts (Criscione, Poulin, & Blouin, 2005; Barrett, Thrall, Burdon, & Linde, 2008; Sweet & Johnson, 2018). Within hybrid zones (HZ) and secondary contact zones (SCZ) this picture is likely to become even more complicated, possibly giving rise to new unexpected patterns. Unfortunately, very few studies have been devoted to this aspect of host-parasite interactions. From the most general point of view, it is assumed that since parasites are dependent on their host, their genetic structure will tend to mirror the host. In this respect, two assumptions are frequently expressed. First, the degree of congruence with the host is dependent on traits connected to the parasitic life-style, such as the degree of host-specificity, transmission mode, presence of dispersal stages, etc. (Maze-Guilmo, Blanchet, McCoy, & Loot, 2016). Generally, the more intimate the association, the higher the degree of congruence. However, this general view may be twisted by many specific traits of the particular host-parasite association. For example, the population structure of heteroxenous parasites (parasites with more than one host in their life cycles) is likely to reflect the least structured host, since any potential structure is erased by the more motile host (Jarne & Theron, 2001; Louhi, Karvonen, Rellstab, & Jokela, 2010). Similarly, with longer free living stage(s), the genetic structures of the host and the parasite become more incongruent (Jarne & Theron, 2001). The second assumption is about the speed of diversification: since the parasites have a shorter generation time, they undergo faster genetic diversification, which may eventually lead to the parasite’s duplication (Page, Lee, Becher, Griffiths, & Clayton, 1998; scenario *a* in Fig. 1). The assumption about the higher mutation rate in parasites was demonstrated in several studies (Nieberding, Morand, Libois, & Michaux, 2004; for ectoparasites: McCoy et al., 2005; Whiteman, Kimball, & Parker, 2007; but see Gómez-Díaz, González-Solís, Peinado, & Page, 2007; Jones & Britten, 2010 for the opposite results).

**Figure 1.**
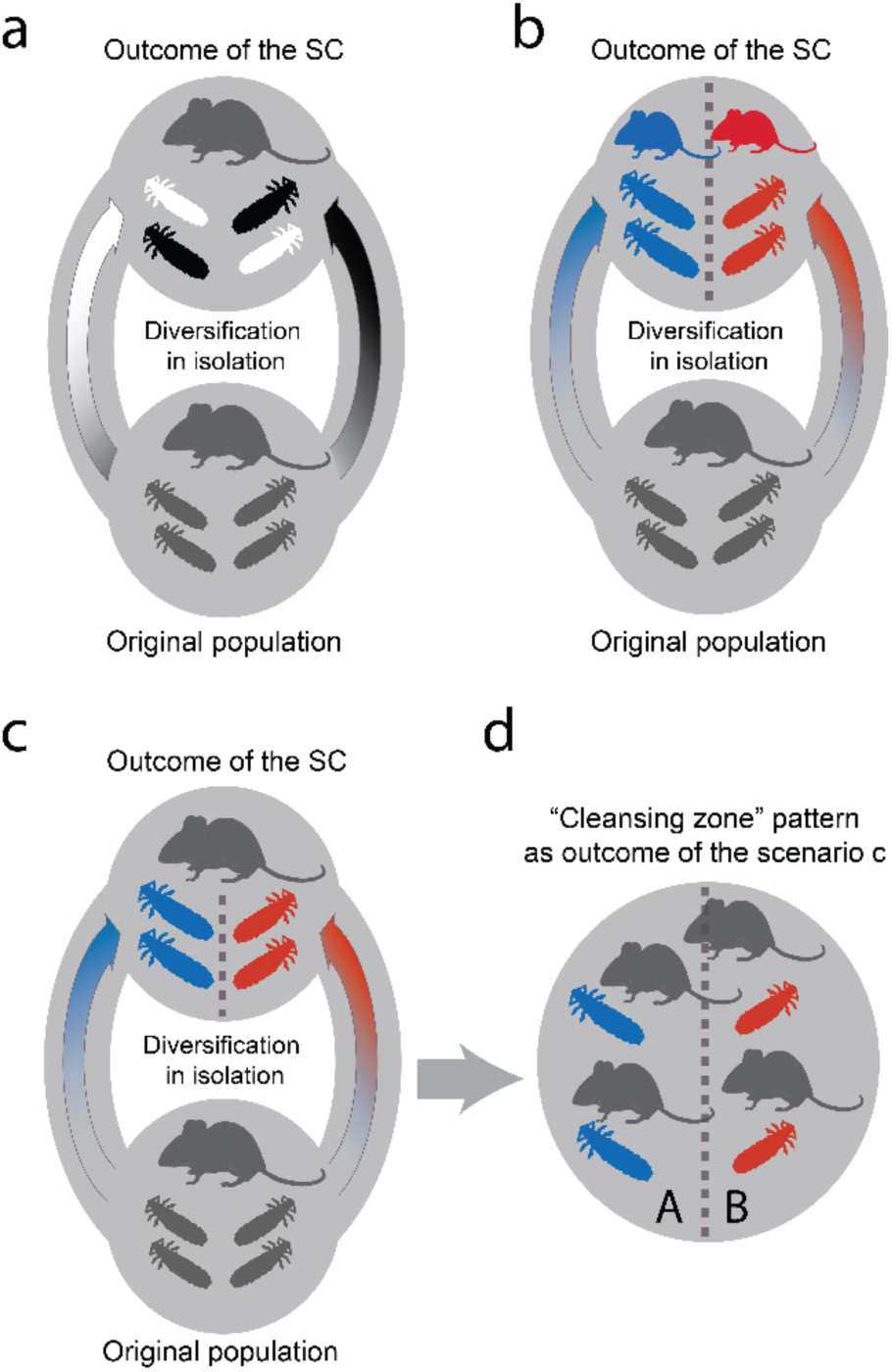
Different scenarios of the secondary contact outcomes in a host-parasite association. **(a)** the host re-establishes a panmictic population while the parasite differentiates into two separated species, which keep spreading and eventually live in sympatry. **(b)** both host and the parasite create HZ; parasite’s HZ is narrower. **(c)** the host re-establishes panmictic population while the parasite creates a narrow HZ. **(d)** as a consequence of the scenario c, the host moves freely across the whole area, while parasite’s lineages are “filtered” by the HZ.

Although many studies have been devoted to comparing phylogenies and population structures of host-parasite associations, only a few analyzed these processes in connection to SCZ and HZ of the hosts (reviewed by Theodosopoulos, Hund, & Taylor, 2019), and only recently, de Bellocq et al. (2018) focused on detecting HZ in parasite populations. Using two parasites of the house mouse *Mus musculus*, the nematode *Syphacia obvelata* and the fungus *Pneumocystis murina*, they found that within the host’s HZ both parasites create their own HZ. They also demonstrated that the parasites (reaching higher genetic divergence) created significantly narrower HZs than the host (scenario *b* in the Fig. 1).

From a theoretical point of view, the assumptions and empirical evidence discussed lead to a third possible scenario: during secondary contact the host does not create a HZ but rather re-establishes a panmictic population, while the parasite accumulates a degree of genetic differences which prevent re-establishment of panmixia but does not lead to speciation (scenario *c* in the Fig. 1). A paradoxical result of such an event would be establishment of a parasite’s HZ within the host’s panmictic population, which on a microevolutionary scale would function as a “host-cleansing filter”: passing through this filter from area A to area B (Fig. 1d), the hosts rid themselves of area A parasites and acquire the area B parasites. To our knowledge, this “filter” has never been observed in nature. In fact, the presence of a parasite’s HZ in the scenario described here is difficult to guess *a priori*, as it is not indicated by the host’s HZ. However, in our previous work (Martinů, Hypša, & Štefka, 2018) we presented the genetic structure of postglacial Europe recolonization by the mice of the genus *Apodemus* and their ectoparasite, the louse *Polyplax serrata*, which corresponds to such scenario (Fig. 2; see below for details).

**Figure 2.**
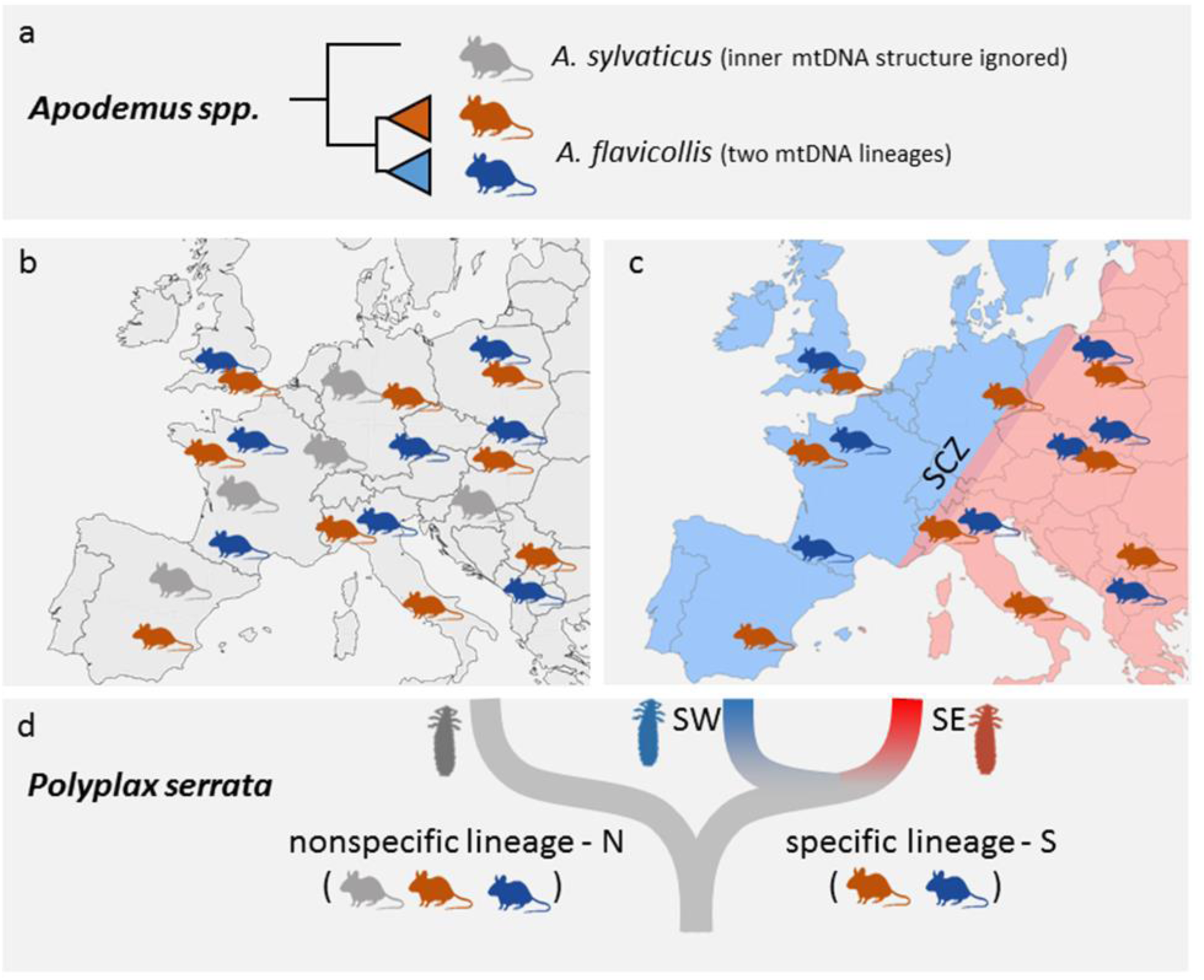
Genetic background of *Apodemus* spp. and *Polyplax serrata* adopted from Martinů et al. (2018). **(a)** The main genetic pattern of *Apodemus* hosts relevant to this study. Inner mtDNA structure of *A*.*sylvaticus* is ignored since not relevant to the discussion of the SCZ (see the text). **(b)** Geographic distribution of the hosts: both spp. co-occur across Europe with their intraspecific mtDNA clades randomly dispersed. Both spp. are parasitized by the non-specific lineage (N) of the lice shown on the d. **(c)** Two mtDNA lineages of *A*. *flavicollis* which in addition to the N lineage are parasitized by the specific lineage (S; deepicted on d). SCZ: schematic representation of the presumed secondary contact zone of the two mtDNA lineages of both the *A*. *flavicollis* and the S lineage of the parasite. While the host’s lineages dispersed across the whole Europe after glaciation, the parasite’s lineages remain confined to the two exclusive areas (red and blue) with a narrow hybrid zone (the violet line). **(d)** The main genetic pattern of *Polyplax serrata* showing ecological complexity of this parasite: the N and S lineages live in sympatry but differ in the degree of host specificity (single host vs two hosts); the SW (specific west) and SE (specific east) lineages create hybrid zone (the „cleansing zone”) which transverses panmictic population of the host.

Similar to all sucking lice, *P*. *serrata* is a permanent homoxenous ectoparasite with strict host specificity, which is transmitted almost exclusively during physical contact of its hosts. As such it falls into the category of highly intimate parasites displaying a high degree of congruence with their hosts. In Fig. 2, we summarize the main features of the population genetic pattern obtained by analysis of 379 bp mitochondrial haplotypes (Martinů et al., 2018). It shows that *P*. *serrata* is composed of several genetic lineages (Fig. 2d) with different host-specificities and geographic distributions. This indicates that even such traits as the degree of host specificity may be very flexible and change rapidly at a shallow phylogenetic level. For example, the so-called *specific* (S) and *non-specific* (N) lineages, although closely related (sister lineages) and living in sympatry, differ in degree of their specificities, one being exclusive to *Apodemus flavicollis*, while the other can also live on *A*. *sylvaticus*. However, the most intriguing part of the pattern was detected within the S lineage. On the mtDNA based phylogenetic trees, the host (*A*. *flavicollis*) and the parasite (S-lineage of *P*. *serrata*), display the same basic structure. Their samples collected across all of Europe form two genetically distant clusters, suggesting recolonization from two different refugia (the taxa designated by red and blue colours in the Fig. 2; see Martinů et al., 2018 for discussion). However, while the two host’s clusters have already spread across the entirety of Europe, their lice did not follow the same process. Instead, their two sub-lineages, designated as *specific east* (SE) and *specific west* (SW), ceased their dispersion after reaching the SCZ in the middle of Europe (Fig. 2). This disparity is surprising since the high intimacy of lice should predetermine them to mirror genetic structure of the host (e.g. Harper, Spradling, Demastes, & Calhoun, 2015).

To obtain a more complex picture of secondary contact in *P*. *serrata*, in this study we analyze three patterns derived from metagenomic data of 13 louse specimens collected across the SCZ: nuclear SNPs, complete mitochondrial genomes, and complete genomes of the symbiotic bacterium *Legionella polyplacis*. We use these analyses to retrieve two kinds of information. First, we compare nuclear and maternally inherited markers (mitochondrial genomes, symbiotic genomes) to demonstrate a narrow HZ between the SW and SE lineages of the lice. Second, we address two possible causes of the SE/SW incompatibility suggested in our previous work (Martinů et al., 2018). *P*. *serrata* carries the intracellular obligate symbiont *Legionella polyplacis* (Říhová, Nováková, Husník, & Hypša, 2017) which could be incompatible with the non-native genetic background. Similarly, since the *Polyplax* louse mitochondria are fragmented into 11 minichromosomes (Dong, Song, Jin, Guo, & Shao, 2014) a rearrangement of their genetic composition could theoretically lead to the SE/SW incompatibility. We therefore compare complete mitochondrial and symbiotic genomes to assess the degree of their divergence.

## Methods

### Origin of the samples

*Apodemus flavicollis* mice were captured across the HZ of *Polyplax serrata*-specific lineages in 2018 in the north-west of Czech Republic, using wooden snap traps. Field studies were carried out with permits approved by the Committee on the Ethics of Animal Experiments of the University of South Bohemia, by the Ministry of the Environment of the Czech Republic, and by the Ministry of the Agriculture of the Czech Republic (No. 51304/ENV/14-2981/630/14, MZP/2017/630/854, 22395/2014-MZE-17214). Mice were searched for lice by visual checking and combing. Lice were removed and stored in 100% ethanol in the −20°C. Genomic DNA from whole louse specimens was individually extracted using the Qiagen QIAamp DNA Micro Kit (Qiagen, Valencia, CA, USA). Membership to individual louse lineages (SE, SW or N) was assigned by sequencing a fragment of the mitochondrial cytochrome oxidase subunit I gene (COI, 379 bp) as in Martinů et al (2018). All sequences of the S lineage available from previous work together with the newly gained specimens (Table S1) were collapsed into haplotypes using ALTER (http://www.sing-group.org/ALTER/). Then the specimens were assigned to individual lineages by phylogenetic reconstruction, using maximum likelihood method with 1000 bootstraps (Fig. S1). Model GTR+I+G was used as the best-fit model, selected according to a corrected Akaike information criterion using jModelTest2 v2.1.10 (Darriba, Taboada, Doallo, & Posada, 2012; Guindon & Gascuel, 2003). Sample DBab5 (PolyN) from the N lineage was used as an outgroup. The S louse lineage was found on 11 *A*. *flavicollis* captured across the contact zone. Thirteen lice (7 SE and 6 SW), 1 to maximum 3 from each parasitized host, were selected for genomic re-sequencing (Fig. 3, Table 1). DNA concentrations were quantified with a Qubit 2.0 Fluorometer (Invitrogen, Carlsbad, CA, USA) using High Sensitivity reagents.

**Table 1.**
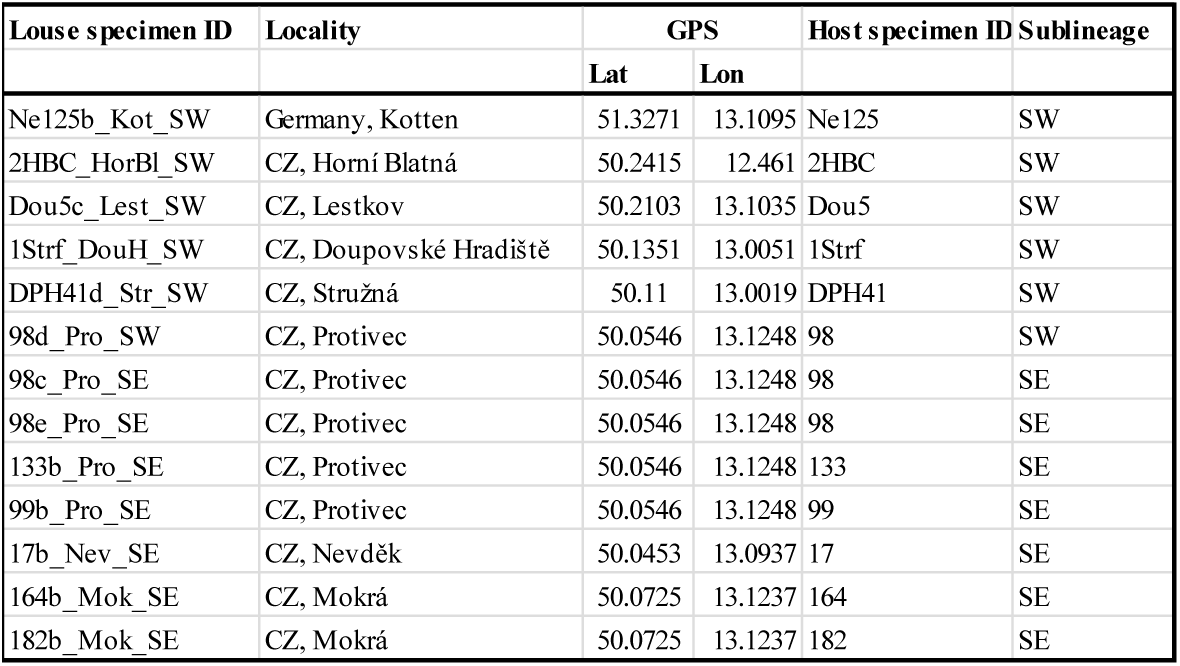
List of the specimens analyzed in this study.

**Figure 3.**
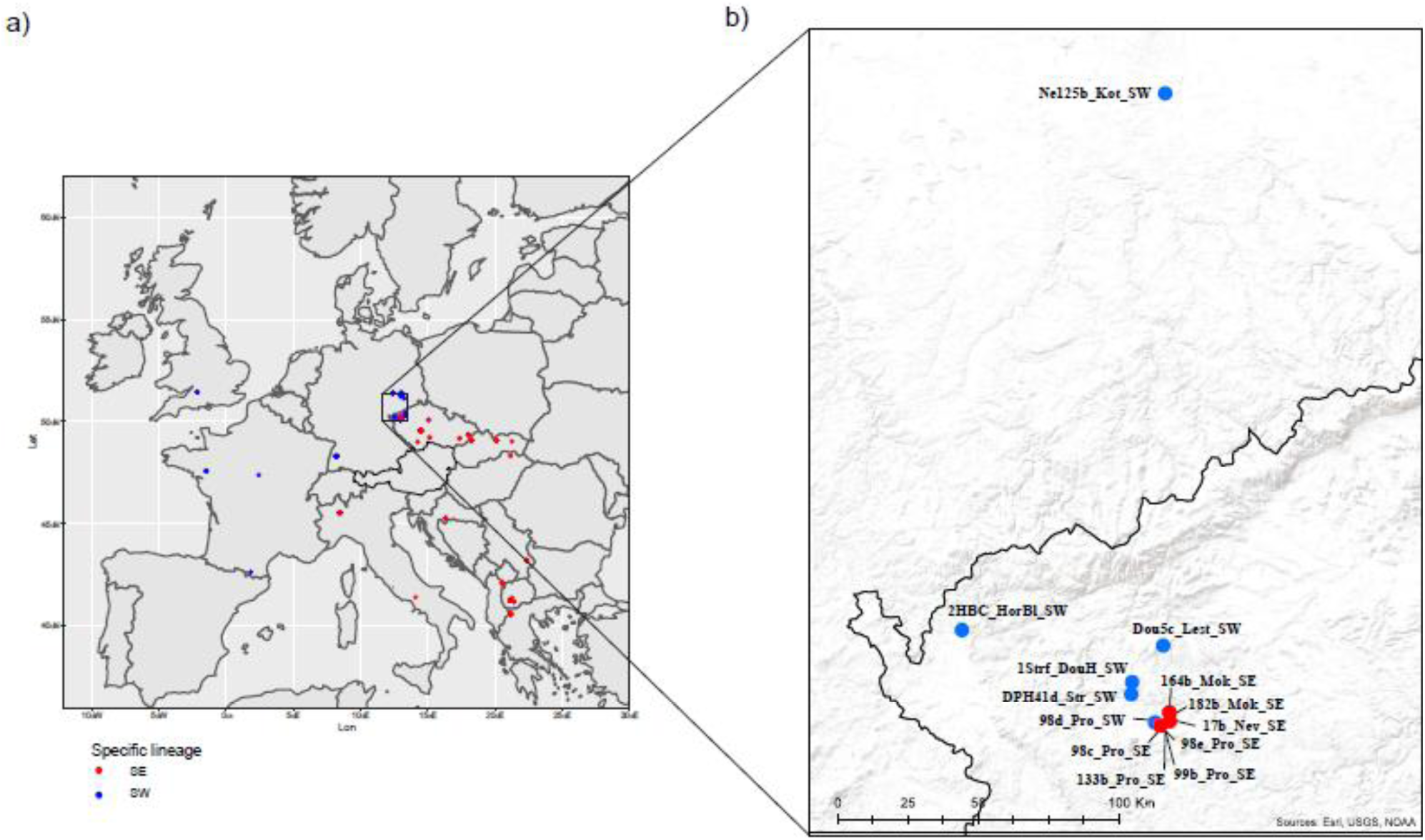
**(a)** Distribution of the *P*. *serrata* S lineage samples analyzed in Martinů et al., (2018). **(b)** Detailed distribution of the samples used in this study. Blue and red designates the SW and SE sublineages, respectively.

### Generating and assembling genomic data

To obtain a reference genome sequence, one *P*. *serrata* specimen (98c_Pro_SE) was sequenced by Oxford Nanopore technology (ONT) on one MinION II flowcell, producing 16.7 Gbp of data (3.3 million reads of average size of 5 kbp). Library preparation and sequencing was performed at Roy J. Carver Biotechnology Center (University of Illinois at Urbana-Champaign, USA). The ONT reads were trimmed with filtlong (v0.2.0; https://github.com/rrwick/Filtlong) to retain sequences of at least 4000 bp and phred score of at least 20. Flye assembler (v2.5; Kolmogorov, Yuan, Lin, & Pevzner, 2019) was used to perform de novo genome assembly using ONT reads (with/without quality trimming) with the expected genome size of 110 Mbp as a parameter. The resulted assemblies were polished once with Racon (v1.3.3; https://github.com/isovic/racon) and twice with Medaka (v0.6.5; https://nanoporetech.github.io/medaka/) using the ONT reads. The final polishing was performed again with Racon, this time using illumina reads. Quality checks of the final polished assemblies were performed with BUSCO (v3; Waterhouse et al., 2017). Biological origin of the assembled contigs was verified by blastx (Altschul et al., 1997) of the polished assembly (98 contigs) against the *Pediculus humanus corporis* genome (GCA_000006295.1 JCVI_LOUSE_1.0). The contigs that did not yield a hit (removed were 60 contigs, 0,46% of the assembly length) were excluded from the following analyses after their non-louse origin was verified by blastx against nr/nt NCBI database.

Whole genome re-sequencing was performed to generate metagenomic data used to map the SNPs, and to assemble genomes of the *L*. *polyplacis* symbionts and mitochondrial minichromosomes. gDNA libraries for thirteen louse specimen were constructed for paired-end Illumina sequencing with NovaSeq6000 instrument. All samples were sequenced on one Illumina Novaseq lane producing on average 62.5 million 150 bp paired-end reads (PE) per sample. Libraries were constructed with an average insert size of 450bp. Fastq files were generated from the sequence data using Casava v.1.8.2 or bcltofastq v.1.8.4 with Illumina 1.9 quality score encoding. All sequencing and fastq file generation was carried out at the W. M. Keck Center (University of Illinois, Urbana, IL, USA). Illumina reads of the re-sequenced specimens were checked for quality and filtered in BBtools (https://jgi.doe.gov/data-andtools/bbtools/), then assembled using the SPAdes assembler v 3.10 (Bankevich et al., 2012), under default settings with the parameter careful, decreasing number of mismatches and indels.

To analyze genetic structure in the area of secondary contact of the *Apodemus*-*Polyplax* populations (and the presumed HZ of the parasite), we compared patterns obtained from three different sources: nuclear SNPs, mitochondrial genomes, and genomes of symbiotic bacterium *L*. *polyplacis*. Our hypothesis (visualized in the Fig.1 c, d) predicts that for both maternally inherited markers (mitochondrial and symbiotic genomes), the inter-lineage divergence (i.e. SE vs. SW) will be much higher than any divergence within the lineages. This reflects the hypothetical scenario of two lineages long isolated during they stay in two different refugia and also during most of the recolonization journey. On the contrary, since the hypothesis presumes rare hybridization, no such clear distinction should be found in nuclear SNPs. To visualize and quantify diversification of maternally inherited markers, we reconstructed genealogical trees and calculated distance matrices for mitochondrial and symbiotic genomes. To visualize nuclear diversification using SNPs, we analyzed population structure using PCA and reconstructed genealogical relationships by building a maximum likelihood tree. Below, we provide details on processing the three sources of the data.

### Nuclear SNPs

Reads from 13 louse genomes were individually mapped with Bowtie2 (Langmead & Salzberg, 2012) to the reference genome described above. Before mapping, reference sequence was indexed using Samtools (Li et al., 2009) and a dictionary file was made with CreateSequenceDictionary in Picard v.2.0.1 (https://broadinstitute.github.io/picard/). After mapping with Bowtie2, resulting SAM files were sorted to the BAM files and indexed using Samtools. Duplicated sequences were removed from sorted BAM files with Picard v.2.0.1 (https://broadinstitute.github.io/picard/) and quality of mapping was verified with QualiMap (http://qualimap.bioinfo.cipf.es/). SNPs for population-level analysis were called using the GATK Genome Analysis Toolkit following the “Best Practices” guide from the Broad Institute (Van der Auwera et al., 2013). SNPs were jointly called for all 13 *Polyplax serrata* samples and filtered with QD (quality by depth) <2.0, FS (Fisher strand test) >60.0, MQ (mapping quality) <40.0, and MQRankSum (mapping quality rank sum test) <–12.5. The SNPs from GATK were filtered with minor allele frequency (MAF) threshold 0.05 in PLINK 1.9 (https://www.cog-genomics.org/plink/1.9/). SNPs in linkage disequilibrium (LD) were pruned with the squared coefficient of correlation (r2) 0.2 and missing data threshold 0.2. Resulting 12, 285 variants passed filters and quality control in 13 *P*. *serrata* samples and used in population structure estimation. Principal Component Analysis (PCA) was performed in R package SNPRelate (DOI: 10.18129/B9.bioc.SNPRelate) and phylogenetic tree based on Maximum likelihood analysis was reconstructed using SNPhylo pipeline (Lee, Guo, Wang, Kim, & Paterson, 2014) with 1000 bootstraps.

### Legionella polyplacis genomes reconstruction and comparison

The contigs corresponding to *Legionella polyplacis* were visually identified using ORF prediction done in the Geneious package (the prokaryotic gene arrangement could be readily recognized by the density of predicted ORFs). In each sample, except for the DPH41, the genome of *L*. *polyplacis* was assembled into a single complete contig. In the DPH41 assembly, we were not able to find the symbiont genome, despite the good assembly quality of the louse genome. Complete *L*. *polyplacis* genomes were annotated in RAST (Aziz et al., 2008) and aligned using Mafft algorithm implemented in Geneious. In three samples (Ne125b_Kot_SW, 99b_Pro_SE, 98c_Pro_SE) the rRNA region did not assemble correctly and was only represented by short fragments. To extend these fragments into full length, we used the program aTRAM 2.0 (Allen, LaFrance, Folk, Johnson, & Guralnick, 2018). Phylogenetic analysis of the resulting 530,042 bp long matrix was performed in Phyml (Guindon et al., 2010). The evolutionary model GTR+I+G was determined for the whole concatenated matrix (considering the strong phylogenetic signal in the matrix, the model selection is a purely formalistic step in the presented analysis) by a corrected Akaike information criterion in jModelTest2 (Darriba et al., 2012) based on the AIC. Comparison of gene content was done manually using the annotated alignment. We were taking into consideration the following criteria: presence of the annotations across all genomes; presence of the corresponding sequence (in case of missing annotation); distribution of differences (i.e. does the difference reflect the SE/SW split).

### Minichromosome genomes reconstruction and comparison

Due to the conserved noncoding region shared by all minichromosomes, the Spades based assembly was not able to separate the different minichromosomes reliably. We therefore took an alternative approach. Using a single gene from each minichromosome as a reference (preliminary assignment of the genes to individual minichromosomes was based on the GenBank data available for *P*. *spinulosa*; acc. nos. KF647762-KF647771) we mapped the reads and extended the sequences in the program aTRAM 2.0 (Allen at al., 2018). To obtain annotations of the resulting sequences, we combined two methods. First method utilized the web based server Mitos (Bernt et al., 2013). Since this method missannotated or entirely missed some of the genes, we used an additional blast based approach. We combined the assembled minichromosome sequences into a custom database and used the 37 genes of *P*. *spinulosa* as blast queries (we run discontinous megablast and tblastx, both with E-value set to 10). We then aligned the *P*. *serrata* minichromosome sequences together with *P*. *asiatica* and *P*. *spinulosa* by Mauve (Darling, Mau, Blattner, & Perna, 2004). These alignments were used as a background for combining and correcting the annotations. To prepare a concatenated matrix, we trimmed all minichromosomes to equal lengths. Phylogenetic tree and genetic distances were retrieved from concatenated alignments by the same approach as for the *L*. *polyplacis* genomes.

## Results

### Gene content and arrangement of the maternally inherited markers

#### Mitochondrial minichromosomes

Neither the Spades nor the aTRAM method was successful in reconstructing complete sequences of the eleven mitochondrial minichromosomes. However, the aTRAM assemblies contained whole coding regions and were used for both the phylogenetic reconstruction and the comparison of gene content between the SE and SW lineages. Although yielding considerable genetic differences (Table S2), the mitochondrial minichromosomes show identical arrangement of the genes (shared synteny) in both the SW and SE lineages (Table S3). This arrangement is also very similar to that in the related louse species, *P*. *spinulosa*. Concatenation of the minichromosomes produced a 17,253 bp long matrix. When phylogenetically analyzed, it yielded a tree with two very distant clusters corresponding to the SE and SW lineages (Fig. 4b, Fig. S2). Within both clusters, the distances were significantly lower than the distance between the clusters. However, the overall range of distance was higher within the SW than within the SE (Table S2, Fig. 4), likely reflecting the broader geographic sampling range of the SW lineage.

**Figure 4.**
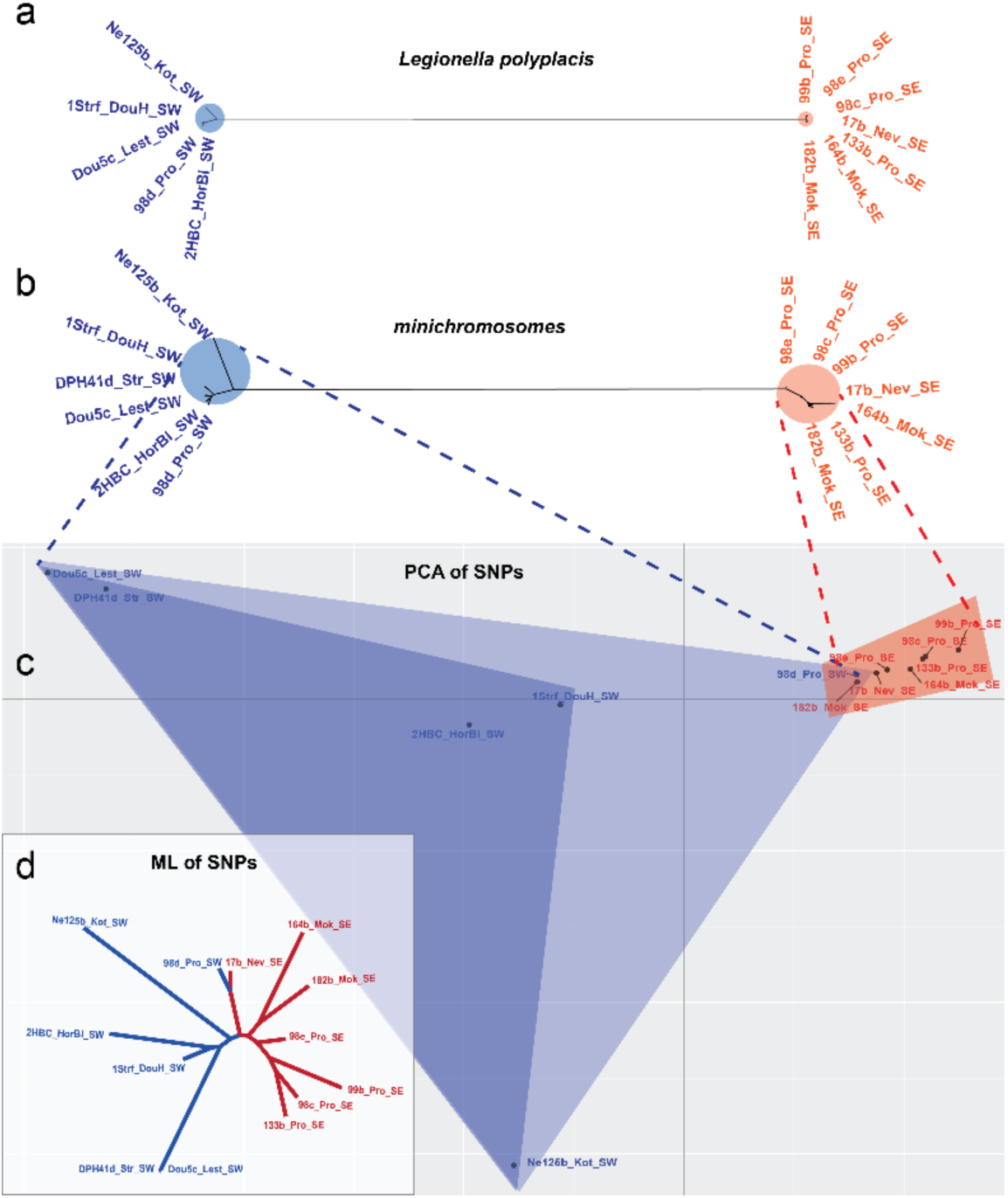
Comparison of maternally inherited markers and nuclear SNPs. In all parts the two lineages are designated by different colours: blue=SW, red=SE. (a) phylogenetic tree based on complete genomes of *L*. *polyplacis* from 12 samples of *P*. *serrata*. (b) phylogenetic tree based on concatenated 11 minichromosomes of *P*. *serrata*. (c) PCA analysis of nuclear SNPs. The light blue triangle designates a complete set of the SW samples, including the 98d_Pro_SW which clusters together with the SE samples (the overlapping red shape); the deep blue triangle designates SW lineage in exclusion of the 98d_Pro_SW sample. (d) ML tree based on the SNPs showing the 98d_Pro_SW sample clustering within the SE clade.

#### Legionella

Genomes of the symbiont *L*. *polyplacis* revealed phylogenomic structure parallel to the mtDNA (Fig. 4a, Fig. S3), with a deep genetic split between the SW and SE lineages. The complete genomes displayed a high degree of similarities with all pairwise comparisons exceeding 99% identity. The contrast between the intra- and inter-cluster comparisons is better illustrated by the counts of the observed differences, which were 27-212 within the SW cluster and 1-57 within the SE cluster, compared to 3,683 - 3,699 between the clusters (Table S2). When comparing the genome sequences, we did not find any clear instance of missing genes. The majority of the gaps introduced by genomic alignment span just one or two nucleotides and were placed in intergenic regions (only one deletion span across 26 nucleotides, also located between the gene coding sequences). The annotations provided by RAST contained several differences between the two clusters, indicating that a gene present in one lineage is shortened or missing in the other cluster. In all of these cases, however, the differences were not caused by a convincing absence of the gene sequence but rather by failure of the algorithm to recognize the sequence as coding a gene, most likely due to the aberrant nature of highly derived symbiont genomes.

#### Diversity and population pattern of nuclear SNPs

Illumina reads from 13 genomes were successfully mapped on the reference with mean coverage ranging from 20,9x (Ne 125b_SW) to 32,1x (98d_Pro_SW). None of the assemblies had to be filtered out due to low coverage and/or low Phred scores. Patterns of population structure obtained using nuclear SNPs differed from genealogies based on maternally inherited genomes (Fig. 4). In contrast to mtDNA, the structure derived by PCA from nuclear SNPs did not recognize two compact and mutually distant clusters. Instead, the distances within the geographically broader SW lineage were comparable to the majority of the SW-SE distances. One of the SW mitochondrial samples (98d_Pro_SW) was significantly closer to the SE samples, and in some analyses even positioned within the SE cluster (Fig. 4, Fig. S4). Within clusters, the distances arranged along the two main axes did not reflect any obvious feature of the samples: for example, the large geographic distance between the NE125b_Kot_SW and other samples was reflected along the main axis, but the distribution of other samples was not strictly correlated with their geography (Fig. 4, see map Fig. 3.).

ML tree-building method yielded a similar pattern to PCA (Fig. 4, Fig. S5). All SE samples clustered together with 98d_Pro_SW sample and created a monophyletic lineage with high bootstrap support. The position of Ne125b_Kot_SW remained unclear, probably due to greater genetic and geographic distance, whereas the rest of SW samples (4 specimens) clustered together.

## Discussion

In this study, we show a new pattern of population structure that can arise at the secondary contact zone (SCZ) due to different courses of evolution in hosts and parasites. The main signature of this pattern is a conflicting arrangement of mitochondrial markers in the host and the parasite at the SCZ (Fig. 2). The host’s mitochondrial lineages, coming from different refugia, mix across the whole area of the host species’ final distribution and re-establish a panmictic population. In contrast, the parasite’s mitochondrial lineages stop their dispersal at the SCZ. In our model, the re-established panmixia of *A*. *flavicollis* is strongly suggested by a previous study on mitochondrial and microsatellite markers (Martinů et al., 2018) and further corroborated by preliminary results of our recent rad-seq analysis (in prep). For the louse *P*. *serrata*, we detected a sharp geographic division between the SE and SW lineages using short (379 bp) cytochrome oxidase I (COI) haplotypes sampled across Europe (Martinů et al., 2018). To obtain a more informative comparison of genetic distance within and between the SE/SW clusters, in the present study we demonstrate this split on near-complete mitochondrial genomes from 13 samples collected across the SCZ (Figs. 3, 4). From a strictly theoretical point of view, the pattern produced by the mitochondrial data can be explained by several scenarios. The first explanation is based on the strong presumption that louse population structure will be determined entirely by the hosts’ migrations, given that the lice are highly host-specific and intimate parasites. Consequently, the discrepancy shown in Fig. 2 would be a sampling or methodological artifact. However, considering the geographic extent and the number of samples in our previous study (Martinů et al., 2018), we believe that a methodological artifact is a highly implausible explanation. This view is further supported by the present study of the complete mitochondrial sequences and the same genealogical pattern obtained for 12 complete genomes of the maternally inherited symbiont *L*. *polyplacis* (Fig. 4).

A second theoretical possibility assumes that the lice speciated during their separation in refugia before secondary contact of their hosts, due to their shorter generation time. A similar case was reported by Hafner et al. (2019) for a recent secondary contact of two subspecies of pocket gophers and their lice. While the gophers established a HZ, their lice had already speciated and their contact resulted in “competitive parapatry”, with one louse species replacing the other. The authors also pointed out that the distribution data on the pocket gophers and their chewing lice indicate many instances of range overlap, potentially representing zones of competitive parapatry or species replacements. There are two strong arguments against applying similar scenarios to our system. A theoretical objection is that since the two *A*. *flavicollis* mtDNA lineages do not create a SCZ or HZ, but intermix across Europe, it is difficult to envisage a mechanism which would prevent dispersion of the two new louse species across the SCZ. Since both louse mtDNA lineages share the same host species and live in identical ecological environments (as evidenced by sampling both lineages even from the same host individuals), their mutually exclusive distribution is obviously not due to different adaptations (i.e. different host/environment specificities). Also, competitive exclusion is a very unlikely cause as demonstrated by the frequent coexistence of the S-lineage and N-lineage (Martinů et al. 2018). An empirical argument rests on the comparison between the mtDNA and SNP data. If the two mitochondrial lineages were fully isolated non-interbreeding species, we would expect to see the same pattern (i.e. two clearly separated and distant clusters) for both the mtDNA and the SNP sets. However, the comparison in Fig. 4 shows that the two sets of data provide very different pictures, where the wide genetic distance between the mitochondrial markers is not matched by similar patterns in the nuclear markers. Although more extensive sampling across the SCZ will be needed to further study the HZ structure in more detail, the visualization based on our 13 SNP sets can be well interpreted. In contrast to the mtDNA, the two SNP sets do not create two distant clusters with the inner genetic diversity negligible when compared to the inter-cluster distance, as would be expected for long-term genetic isolation (Figs. 4, S4, S5). The positions of the individual samples in the PCA plot of the SNPs are determined partly by their geography and partly by their origin within the SW/SE lineage. As a consequence, the SE individuals, sampled at several close localities, form a tight cluster while the SW individuals, sampled from a considerably larger area, are scattered in the PCA plot and their distances are comparable to (or even larger than) the distance between the two clusters. Moreover, the SW sample 98d_Pro_SW collected from the same locality (and same individual mouse) as some of SE samples was genetically closer to the SE population than to any of the SW samples.

The third hypothesis assumes that during their separation, the two parasite lineages reached a high degree of genetic differentiation resulting in a strong but not absolute postzygotic barrier, whilst lacking an efficient prezygotic barrier preventing them from mating. As a consequence, upon encountering each other they formed an extremely narrow HZ in which the majority of the inter-lineage mating’s fail. In this case we would expect a sharp geographic division between the SW and SE population and genetically close or overlapping clusters of the nuclear markers around the SCZ. Based on the data presented in this study and the previous extensive analysis of mtDNA (Martinů et al., 2018), we consider this hypothesis to be the best explanation of the observed patterns. A decoupled genetic structure of a host and its parasite(s) is not exceptional. It has been reported in various host-parasite associations and caused by different biological and/or environmental circumstances (e.g du Toit, van Vuuren, Matthee, & Matthee, 2013; Hafner et al., 2019). However, to our knowledge, the *Apodemus*-*Polyplax* association presented here is the first known example of genetic structuring caused by a parasite’s HZ created in the absence of the reciprocal host’s HZ. There are several possible factors behind the lack of evidence for similar patterns in nature. Firstly, only a few studies have dealt with HZ in parasites, and they were usually approached in relation to their hosts’ HZ (e.g. Theodosopoulos et al. 2019). This is understandable considering the prevailing view of parasites’s evolution being predominantly determined by their hosts. Secondly, it is likely that emergence of this pattern during secondary contact requires “well-tuned” ratios of genetic diversification between host and parasite populations. If the diversification is too strong, it may either result in speciation of both counterparts (i.e. classical cospeciation Page, 2003), in speciation of the parasite and emergence of HZ in the host (Čížková et al. 2018; Hafner et al. 2019) or in HZ for both counterparts (e.g. de Bellocq et al. 2018). On the contrary, if the diversification is too weak, both counterparts will re-establish panmictic populations. This only leaves a narrow window of time for hosts’ panmixia vs. parasite’s HZ. Yet, such cases do not necessarily have to be rare in nature, they may just be understudied or unnoticed due to the *a priori* view that evolution in host-specific parasites is linked to their hosts. The case we present here shows that one possible indication of a decoupled pattern is strong mtDNA structure in a panmictic host population.

Genetic incompatibility between two populations at the SCZ can be caused by various mechanisms. Apart from the differences accumulated in the nuclear genetic information, interbreeding can also be prevented by incompatibility of mitochondrial and nuclear genetic information (Wolff, Ladoukakis, Enríquez, & Dowling, 2014). In our system, the lice are known to have their mitochondrial DNA split into several circular minichromosomes (Cameron, Yoshizawa, Mizukoshi, Whiting, & Johnson, 2011; Song et al., 2019). The distribution of mitochondrial genes among the minichromosomes is not entirely conserved - there are several differences in the gene arrangements when comparing the species *Polyplax spinulosa* and *P*. *asiatica* (Dong et al., 2014). To address the possibility of mitochondrial gene rearrangement as the barrier between the SW and SE lineages, we reconstructed full minichromosomes (their coding part) from all sequenced samples. In all cases, we found the same gene arrangement, indicating that mitochondrion-nucleus incompatibility is probably not causing the SE-SW isolation (Table S3). Although we cannot exclude existing differences in the non-coding part of the minichromosomes, available information suggests a high level of conservation between minichromosomes within and between related species (Dong et al., 2014). Thus, we assume no mito-nuclear incompatibility to occur in the noncoding part either. In a similar way, we were not able to detect any significant difference between the *L*. *polyplacis* genomes from the SE and SW lineages. While, strictly speaking, the high similarity of the SE and SW in respect to the gene content of minichromosomes and symbiotic genomes is not a refutation of the idea that mitochondrial-nuclear and/or symbiont-nucleus incompatibility could cause the barrier, it makes it at least unsubstantiated.

It would be speculative to infer other genetic sources of incompatibility between SE and SW lineages without a detailed study of the louse nuclear genome and more extensive sampling in the SCZ, which is beyond the scope of the current study. Nevertheless, based on evidence collected from three genetic resources, the two maternally inherited markers (*Legionella* and mtDNA) and nuclear SNP diversity, we were able to unambiguously distinguish between the three possible scenarios of host-parasite incongruence. We propose a new mechanism in host-parasite co-evolution, where a narrow HZ is present in the parasite without a corresponding break in the genetic structure of its host. In this way, the panmictic population of the host is “cleaned” of the parasite lineage present on one side of the parasite’s HZ and replaced by a different parasite lineage on the other side. Given that this evolutionary scenario can easily pass unnoticed (due to the lack of structure in the host) we hypothesize that “host-cleaning filters” may be more common than is currently known, particularly in highly host-specific parasites.

## Acknowledgements

We thank colleagues and students at the FSci USB for assistance during field collecting, we thank Jakub Vlček for providing help with adapting several bioinformatic scripts. This work was supported by the Grant Agency of the Czech Republic (grant number 17-19831S to VH). Access to computing and storage facilities owned by parties and projects contributing to the National Grid Infrastructure MetaCentrum provided under the programme “Projects of Large Research, Development, and Innovations Infrastructures” (CESNET LM2015042), is greatly appreciated. We thank Joel J. Brown for his language corrections on this manuscript.

## Author Contributions

J.M. performed laboratory work, J.M. and A.P made data analyses under the supervision of J.Š. and V.H., with V.H. and J.Š. conceiving the study of *P*. *serrata* S lineage secondary contact zone. All four authors contributed toward the design of the study, and drafted the manuscript.

---

**Fig. S1.**
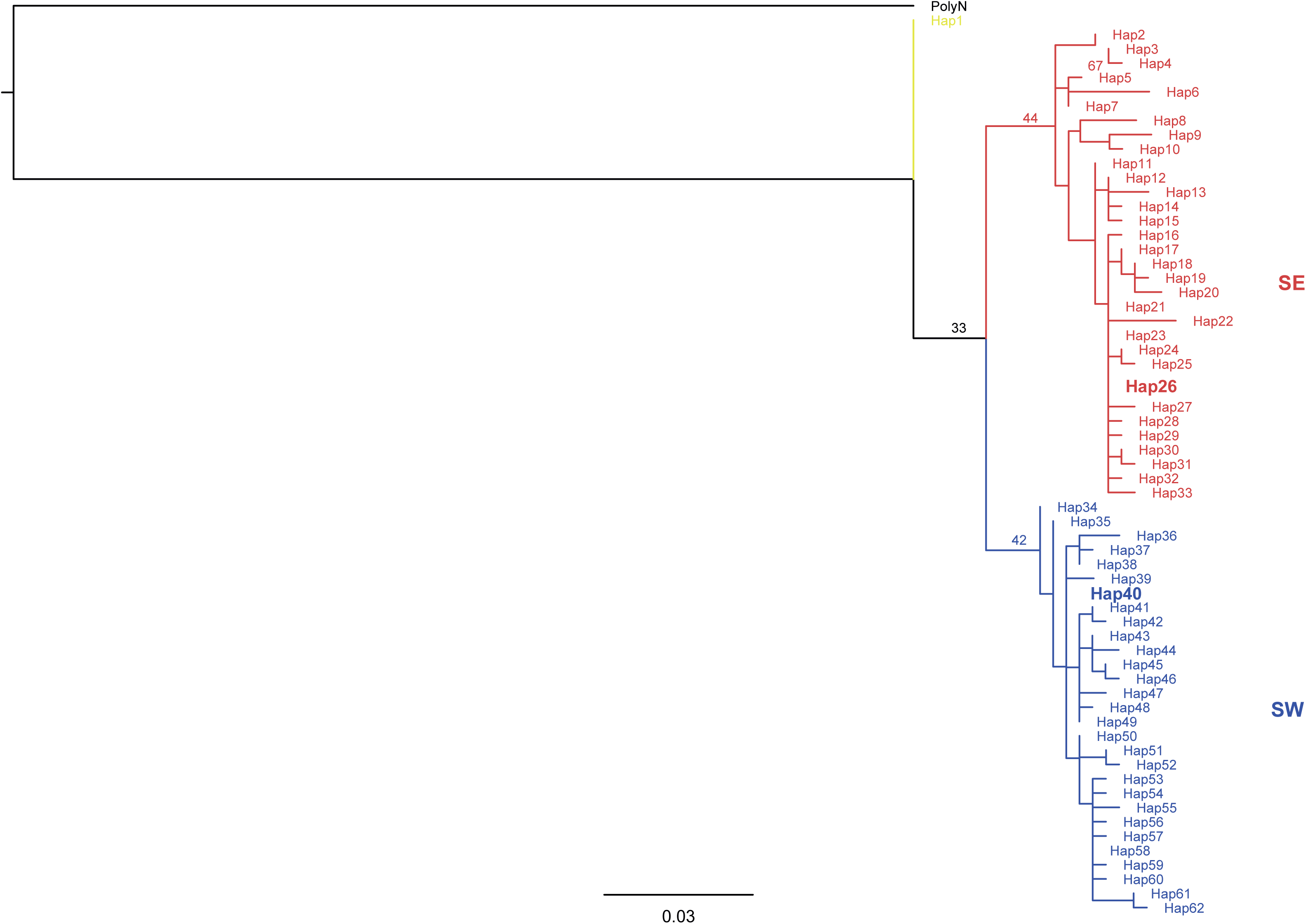
ML tree of the *P*. *serrata* S lineage based on COI haplotypes. It shows the relationship of haplotypes representing 347 samples analyzed in Martinů et al. (2018). Position of the 13 samples analyzed in this study is indicated in bold (the Hap 26 and 40). Blue and red designates the SW and SE sublineages, respectively.

**Fig. S2:**
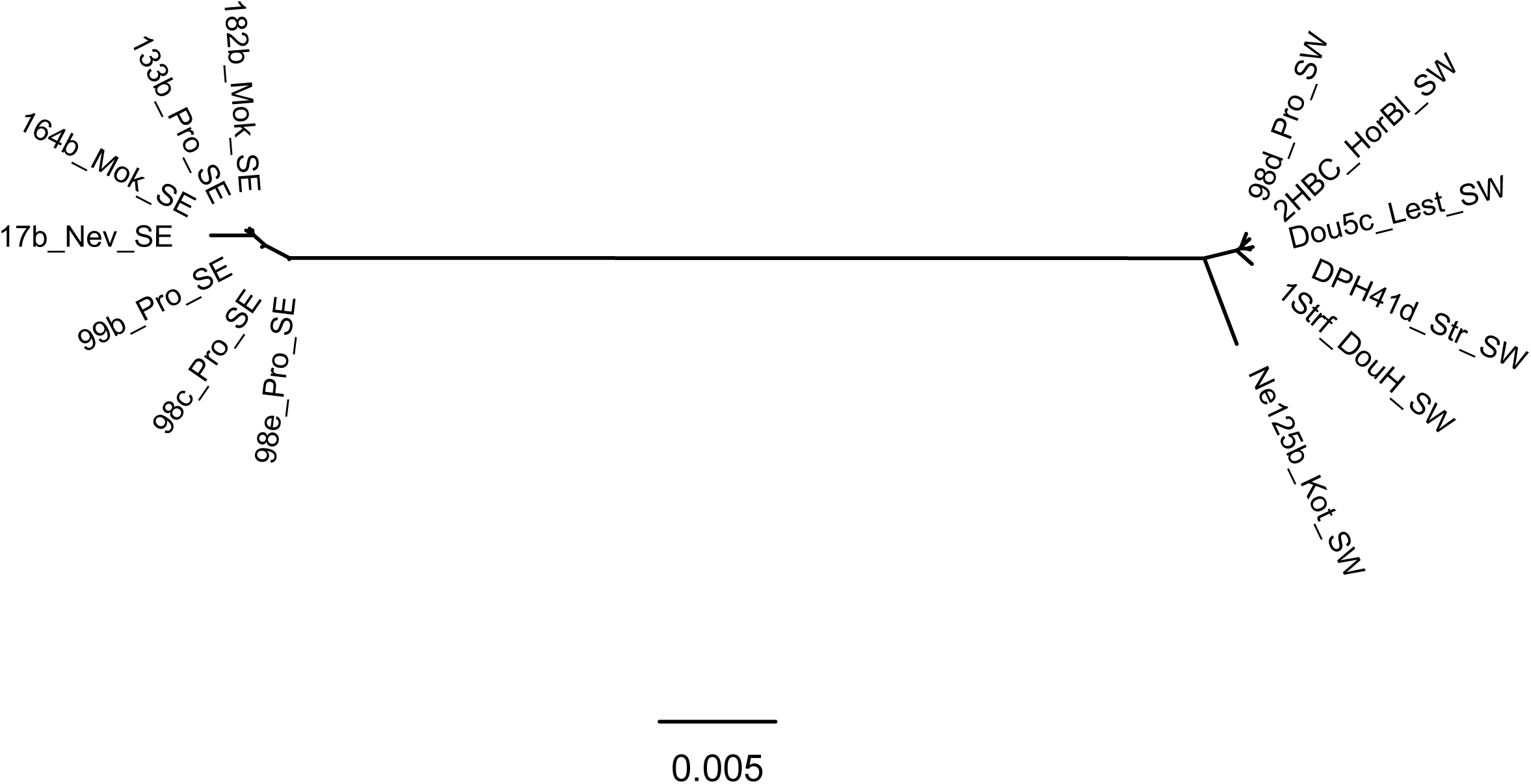
ML tree inferred from concatenated matrix of 11 mitochondrial minichromosomes.

**Fig. S3:**
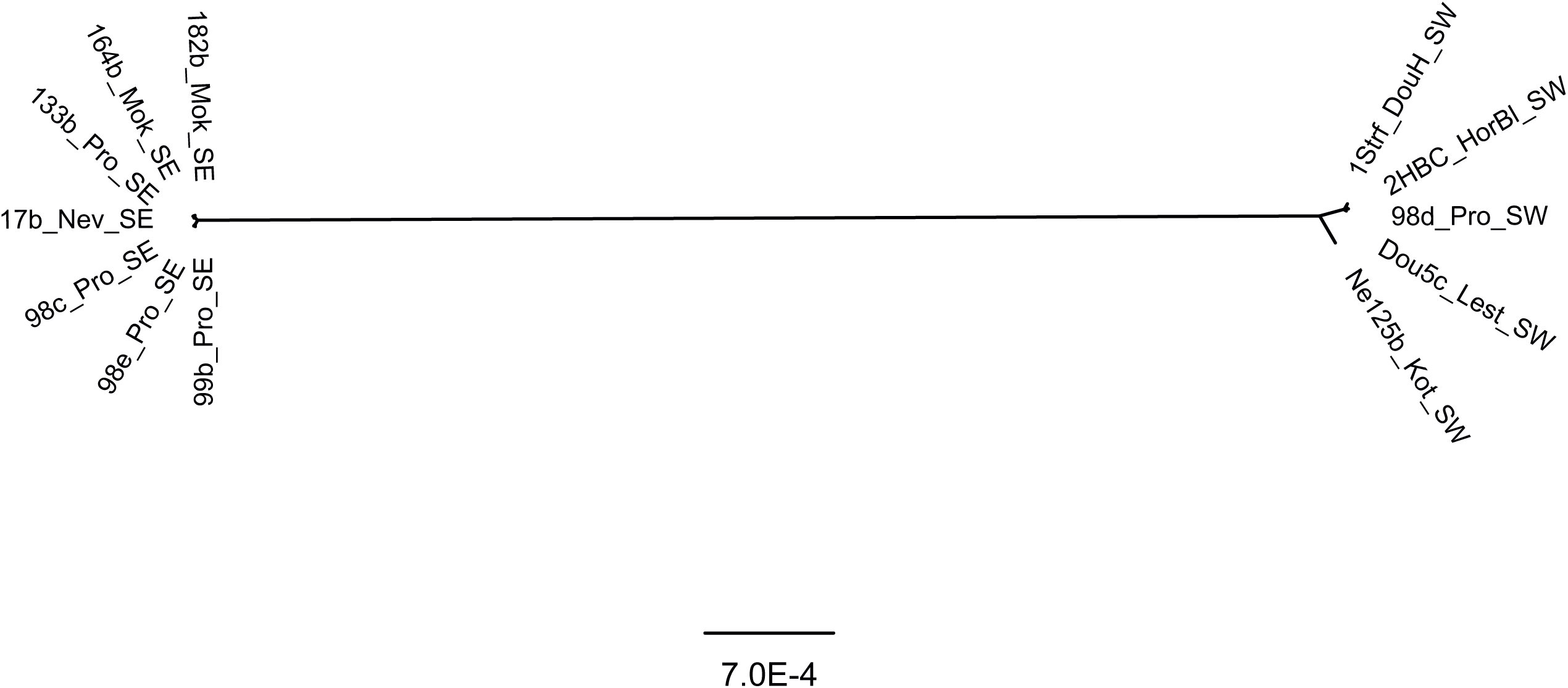
ML tree inferred from complete genomes of the symbiont *Legionella polyplacis*.

**Fig. S4:**
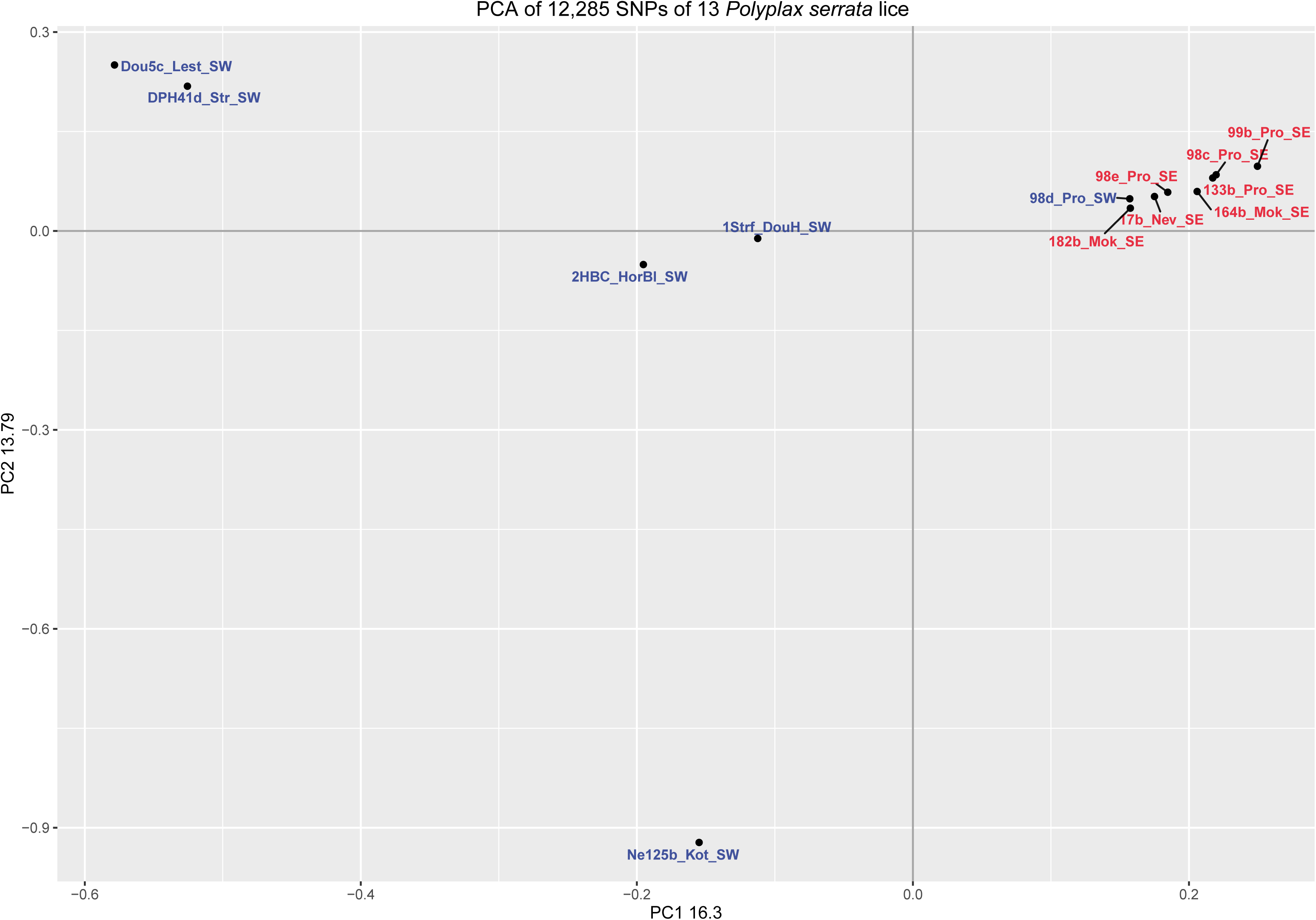
PCA analysis of 12,285 SNPs. Blue and red designates the SW and SE sublineages, respectively.

**Fig. S5:**
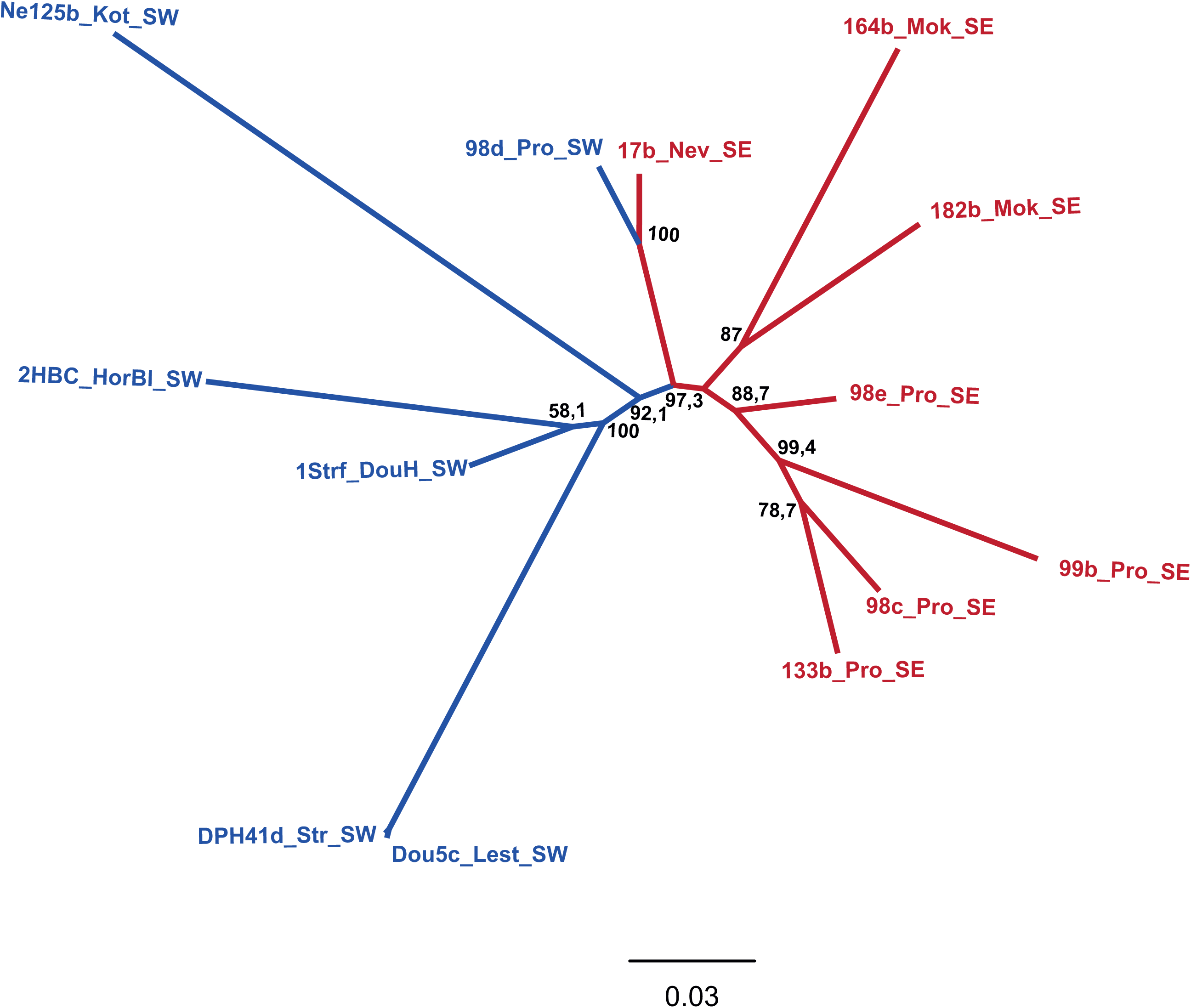
ML tree derived from 12,285 SNPs. Blue and red designates the SW and SE sublineages, respectively.

